# DanioCTC: Injection of circulating tumor cells from metastatic breast cancer patients in zebrafish xenografts for analysis of metastasis

**DOI:** 10.1101/2023.06.05.543673

**Authors:** Florian Reinhardt, Luisa Coen, Mahdi Rivandi, André Franken, Eunike Sawitning Ayu Setyono, Tobias Lindenberg, Jens Eberhardt, Tanja Fehm, Dieter Niederacher, Franziska Knopf, Hans Neubauer

**Affiliations:** Department of Obstetrics and Gynecology, Heinrich Heine University of Duesseldorf, Duesseldorf, Germany; Center for Regenerative Therapies TU Dresden (CRTD), Center for Molecular and Cellular Bioengineering (CMCB), TU Dresden, Dresden, Germany; Center for Healthy Aging, Faculty of Medicine Carl Gustav Carus, TU Dresden, Dresden, Germany; Anatomical Institute, Neuroanatomy, Medical Faculty, University of Bonn, Bonn, Germany; ALS Automated Lab Solutions GmbH, Jena, Deutschland

**Keywords:** breast cancer, circulating tumor cells, zebrafish, in vivo model, metastasis

## Abstract

Circulating tumor cells (CTCs) are considered as metastatic precursor cells, and zebrafish xenografts provide an *in vivo* model to study cancer cell spread. Currently, the low number of patient-derived CTCs limits their analysis in animal models. We present DanioCTC, a xenograft workflow for injecting CTCs from metastatic breast cancer (MBC) patients into zebrafish embryos to study cell dissemination *in vivo*. The study successfully adapts existing workflows and combines diagnostic leukapheresis (DLA), the Parsortix microfluidic system, flow cytometry, and the automated cell micromanipulator CellCelector setup to enrich and isolate MBC-derived CTCs and to finally inject them into Zebrafish embryos, where their dissemination was tracked up to 3 days post-injection. MDA-MB-231 cells were used as a standard xenotransplantation control, and these cells were frequently found in the head and blood-forming regions of the tail. Using DLA aliquots spiked with MBA-MB-231 cells, the newly established DanioCTC workflow confirmed the dissemination of MDA-MB-231 cells into these regions. CTCs from an MBC patient were then enriched by DLA, Parsortix, and flow cytometry, isolated with the CellCelectorTM and xenografted into zebrafish embryos. CTCs were mainly detected in the head and trunk, unlike MDA-MB-231 cells, which were present in the head and tail. DanioCTC presents a significant breakthrough in the use of zebrafish embryos as a model to study CTC dissemination *in vivo*, which can be used for patient-derived CTCs instead of cell culture-derived cancer cells as a crucial step towards understanding the biology of metastatic breast cancer.

**Statement of significance:** DanioCTC is a novel workflow to inject patient-derived CTCs into zebrafish, enabling studies on CTC dissemination and personalized treatment *in vivo*, therefore advancing our toolkit to fight metastatic cancer.

## INTRODUCTION

Breast cancer metastasis is a complex process involving tumor cell migration, intravasation, survival in the bloodstream, adhesion, extravasation, and colonization at secondary sites (1, 2). Metastatic breast cancer (MBC) is the major cause of breast cancer morbidity and mortality, and identifying the mechanisms underlying the metastatic process and developing clinical strategies to detect and treat MBC patients is crucial (3).

Circulating tumor cells (CTCs) are shed into the blood by tumor tissue and are considered precursor cells for metastasis formation (4). Despite being extremely rare, CTC numbers correlate with poor survival outcomes in metastatic and non-metastatic cancers, making their isolation and analysis important for therapy prediction and for understanding their function (5-9). CTC analysis also allows monitoring of the treatment response (10).

One technology to increase the number of informative cancer patient samples containing high numbers of analyzable CTCs is diagnostic leukapheresis (DLA), a density-based blood separation that facilitates the safe collection of large CTC numbers from liters of patient blood (11). DLA enables CTC-positivity rates of more than 90% and a 30-fold increase in CTC-numbers compared to normal blood volumes (7, 12, 13). Moreover, DLA-isolated CTCs are viable and can be used in *in vitro* assays or for the development of preclinical animal models (14), both of which are essential for investigating the metastatic biology of CTCs and for initiating novel effective treatment strategies.

The limitations of current mouse xenotransplantation models to study early tumor dissemination have prompted the need for alternative animal models. Zebrafish have emerged as a valuable tool for investigating human diseases, including cancer metastasis, owing to their genetic similarity to humans, ease of maintenance, and high fecundity (15-17). Zebrafish embryos and larvae are transparent, allowing for non-invasive experimental procedures such as high-resolution *in vivo* microscopy, which provides insights into cancer cell distribution, invasion, and proliferation (16, 18). Although zebrafish xenotransplantation models have been developed to study various malignancies (19), the experimental requirements of microinjection for xenotransplantation currently rely on high tumor cell numbers and/or (transient) *in vitro* cultivation, which limits the use of zebrafish (as other animal models) in studying CTCs (20, 21). To overcome this limitation and to expand the repertoire of xenotransplantation approaches in zebrafish, we have developed a novel workflow for injecting a few isolated CTCs from MBC patients into zebrafish embryos. This approach provides a clinically relevant setting to investigate the CTCs’ metastatic potential in zebrafish, paving the way for personalized therapy approaches and the discovery of new therapeutic strategies for metastatic cancer treatment.

## METHODS

### Cell line and culture conditions

MDA-MB-231 cells were obtained from the American Type Culture Collection (Manassas, Virginia); EGFP-labeled MDA-MB-231 cells were kindly provided by Manja Wobus/Martin Bornhäuser. MDA-MB-231 cells were cultured in RPMI 1640 medium, containing 10% fetal bovine serum, 25 mM HEPES (Gibco) and 100 units/ml penicillin-streptomycin in a humidified incubator at 37°C with 5% CO2; EGFP-labeled MDA-MB-231 cells were cultured in low glucose DMEM (1 g/ml D-Glucose, Gibco) with 10% fetal bovine serum (Gibco) in a humidified incubator at 37°C with 5% CO2. Cultured cells were harvested at a confluence of approximately 80% for staining and spike in experiments.

### Zebrafish and housing conditions

The zebrafish (*Danio rerio*) strains were maintained according to national law and under standardized conditions as previously described (22, 23). Lines used in this study were Tg(*kdrl*:EGFP) (24) kindly provided by Dr. Bernhard Fuss/Prof. Dr. Didier Stainier and Tg(*osx*:mCherry) [Tg(*OlSp7*:mCherry)zf131] for initial characterization of MDA-MB-231 cancer cell distribution in zebrafish embryos (25).

### Standard injection of cell line cells into zebrafish embryos

EGFP labeled MDA-MB-231 cells were trypsinized using trypsin–EDTA (0.25%), neutralized using complete medium, centrifuged at 1,200 rpm for 10 min, re-suspended in PBS, and additionally stained with SP-DiOC18(3) (D7778, Invitrogen) according to the manufacturer’s recommendations. The stained cell suspension was centrifuged for 5 min at 1,200 rpm and the cell pellet suspended in transplantation buffer [0.9x PBS, 1.5 mM EDTA (Roth), 1% Penicillin-Streptomycin (Roth)] at 100–150 cells/nl. Glass needles for injection were pulled from glass capillaries with filament (TW100F-3, WPI) with a P-97 Flaming/Brown micropipette puller (Sutter Instrument Co.). Zebrafish embryos (2 days post fertilization, dpf) were manually dechorionated and anesthetized using 0.02% tricaine and transferred to a petri dish containing 1.5% low melting agarose in E3 (50 mM Natriumchlorid, 0.17 mM Kaliumchlorid, 0.33 mM Calciumchlorid, 0.33 mM Magnesiumsulfat). The labeled MDA-MB-231 cell suspension was loaded into a pulled glass capillary and 1-2 nl were micro-injected into the blood circulation of Tg(*osx*:mCherry) zebrafish embryos via the duct of Cuvier (DoC) using a pneumatic PicoPump (SYS-PV820, WPI, Sarasota, USA). Engrafted embryos were maintained in a new petri dish at 33 °C (26). Based on the fluorescence spread of the injected embryos post injection, embryos with tumor cells in the blood circulation were selected for maintenance for up to three days post injection (dpi). Anesthetized zebrafish embryos were imaged daily starting on the day of injection with an Olympus MVX10 microscope to inspect survival, cardiac edema, homing of tumor cells and extravasation events. The imaging time was kept to a minimum to ensure minimal exposure to anesthesia. No effect on developing larvae was observed during the experimental period, except for one larva which did not survive up to 3 dpi and which was therefore excluded from the analysis (n = 6 throughout the experiment). Data of 1 dpi and 3 dpi are shown. For analysis, the larval body was divided into three regions: head, trunk, and tail (Supplement Figure 1). Quantification of absolute numbers of cell dissemination were performed using the cell counter feature in Fiji.

### Workflow for injection of CTCs into zebrafish embryos

#### Diagnostic Leukapheresis

Diagnostic Leukapheresis (DLA) was performed at the Department of Transplantation Diagnostics and Cell Therapeutics, Duesseldorf, Germany, as previously described (11, 13). Aliquots of the DLA sample were frozen and stored for future use. Cryopreservation of DLA samples was described by our group before (14).

#### Enrichment of viable CTCs or spiked cell line cells from DLA samples with the Parsortix system

Cryopreserved DLA samples were rapidly thawed in a 37°C water bath and filtered through a fine sieve (mesh size 100 μm). The filtered DLA sample was diluted 1:20 in PBS. For spike in experiments, MDA-MB-231 cell line cells were spiked into diluted negative control DLA samples. Therefore, cultured MDA-MB-231 cells were trypsinized using trypsin–EDTA (0.05%), neutralized using complete medium, centrifuged at 1,200 rpm for 5 min and re-suspended in PBS. Subsequently, 5,000 MDA-MB-231 cell line cells were transferred into diluted negative control DLA samples. Viable CTCs of an MBC DLA sample or spiked cell line cells of negative control DLA samples were enriched with the FDA approved Parsortix system (Angle plc). The Parsortix system was used according to the manufacturer’s instructions. Following the protocol, filtration cassettes with 6.5 μm gaps were used and 100 mbar of pressure was applied (14, 27).

#### Isolation of cell line cells and CTCs by flow cytometry and staining for cell tracking

Parsortix pre-enriched MDA-MB231 cell line cells or CTCs were further isolated by flow cytometry. Therefore, enriched MDA-MB-231 cell line cells or CTCs were stained for EpCAM (1:50, VU1D9, Stemcell Technologies, Vancouver, Canada, AF488) and CD45 (1:25, 35-Z6, Santa-Cruz Biotechnology, Dallas, USA, AF647) at 37°C for 60 min for identification by flow cytometry. Cell suspensions were washed twice with PBS at 1,500 rpm for 4 min and were resuspended in PBS. The protocol used for flow cytometry consisted of a discrimination of FITCpos/neg and CD45pos/neg events. FITCpos/CD45neg cells were sorted into a 1.5 ml tube. Subsequently, cells were additionally stained with 10 μM CellTracker Red CMPTX (Cat. No. C34552, Thermo Fisher Scientific) at 37°C for 30 min for cell tracking in zebrafish larvae. Cell suspensions were washed twice with PBS at 1,500 rpm for 4 min and resuspended in transplantation buffer.

#### Detection and picking of single CTCs and cell line cells using the CellCelector

Detection and isolation of single stained CTCs and MDA-MB-231 cell line cells was performed with the CellCelector (ALS GmbH, Jena, Germany)(28), which is a semi-automated micromanipulator consisting of an inverted fluorescent microscope (CKX41, Olympus, Tokyo, Japan) with a CCD camera system (XM10-IR, Olympus, Tokyo, Japan) and a robotic arm equipped with a vertical glass capillary. A 20 μm capillary (CC0048 ALS GmbH, Jena, Germany) with a broken tip was used for cell isolation as well as injection. Cell suspensions were transferred onto glass slides, placed on the automatic stage of the CellCelector microscope and were allowed to settle. Samples were manually scanned with 20× and 40× magnification using the following channels: brightfield (BF, cell morphology), FITC (EpCAM), TRITC (CellTracker) and Cy5 (CD45). The following exposure times were used: 200 ms (FITC), and 200 ms (Cy5). 50 single cells were picked into the capillary meeting the criteria round, intact morphology in BF, FITC positive/TRITC positive/Cy5 negative. The cells were allowed to settle in the capillary tip for 30 min before injection into the zebrafish embryos in order to minimize injection volume.

#### Injection of stained CTCs and cell line cells into zebrafish embryos by combining the CellCelector with a stereomicroscope

Transgenic *Tg*(*kdrl:EGFP*) zebrafish embryos (2 dpf) were manually dechorionated and anesthetized using 0.02% tricaine and transferred to a petri dish containing 1.5% low melting agarose in E3. The petri dish with the transgenic *Tg*(*kdrl:EGFP*) zebrafish embryos was positioned in the deposit area of the CellCelector. An additional Nexius Zoom EVO stereomicroscope with a flexible arm was installed to visualize the embryos in the deposit area of the CellCelector. Approximately 5 – 10 nl of the capillary volume were semi-automatically micro-injected with the CellCelector into the DoC of multiple zebrafish. Engrafted embryos were maintained in a new petri dish at 34°C. Based on the fluorescence spread of the injected embryos at 2 h post injection (hpi), embryos with tumor cells in the blood circulation were selected for maintenance for up to 3 dpi. Zebrafish embryos were imaged at 1 dpi and 3 dpi with the fluorescent microscope of the CellCelector to inspect survival, cardiac edema, homing of tumor cells and extravasation events. The imaging time was kept to a minimum to ensure minimal exposure to anesthesia. No effect on developing larvae was observed during the experimental period.

The complete duration of the workflow – from CTC enumeration to zebrafish CTC injection - was 3 - 4 h.

#### Patient samples

Patient samples were selected from the Augusta study (approved by the Ethics Committee of the Medical Faculty of the Heinrich Heine University Düsseldorf; Ref-No: 3430). Analysis of human samples was carried out in accordance with the Good Clinical Practice guidelines. All patients provided written informed consent for the use of their blood samples for CTC analysis and for translational research projects. Injected CTCs were isolated of a 57-year-old patient with a hormone-receptor positive, Her2/neu negative breast cancer with bone, bone narrow and lymph node metastases.

### Statistics

Statistical analysis was performed using the Prism analysis program (GraphPad Inc., San Diego, CA, USA). Data are expressed as mean ± SD. Statistical analyses were performed by a Repeated Measures ANOVA followed by Bonferroni’s tests or by a Friedman test with Dunn’s post hoc test. p < 0.05 was considered to be statistically significant (*0.01 < p < 0.05; **0.001 < p < 0.01; ***0.0001 < p < 0.001).

### Data Availability

The data generated in this study are available upon request from the corresponding authors.

## RESULTS

### Development of a single cell injection workflow - concept and challenges

We set up a single cell injection workflow by combining, adapting and fully integrating existing, well-validated workflows for each of the required technology modules: (i) xenotransplantation of cancer cells into zebrafish embryos by microinjection into the blood circulation; (ii) DLA, which allows to process large volumes of blood for CTC analysis (iii) flow cytometry for the provision of well-characterized CTC preparations; and (iv) the CellCelector automated micromanipulation system – extended by an additional optical system – for isolation of single cells and performing their injection. It is worth mentioning that - although injection of cells into zebrafish is an already well-established method, high cell number preparations are currently required for xenotransplantation, restricting their use to cell lines or transiently cultured primary cells. The main challenge for the establishment of the presented DanioCTC approach was to fine-control the injection process and to optimize the design of the glass capillary used for picking and injecting selected CTCs at desirable injection rates while maintaining acceptable health conditions for the embryos. In addition, live cell labelling and fluorochrome detection for tracking the injected cells in the living zebrafish embryo were improved, as CTCs, due to challenges in cultivating them, cannot be efficiently transfected, in contrast to cell lines. In addition, direct injection of CTCs without any preculture is preferred to inspect their function during metastasis.

We followed the above concept by first using the invasive, triple-negative MDA-MB-231 cell line to establish the microinjection procedure and to test for the cells’ distribution and survival in zebrafish embryos. This was followed by the development of the entire DanioCTC workflow including the cell line’s live cell labelling and isolation from spiked-in DLA samples for their *in situ* tracking. Ultimately, the workflow was adapted and used to inject and monitor patient derived CTCs (Figure 1).

**Figure 1:**
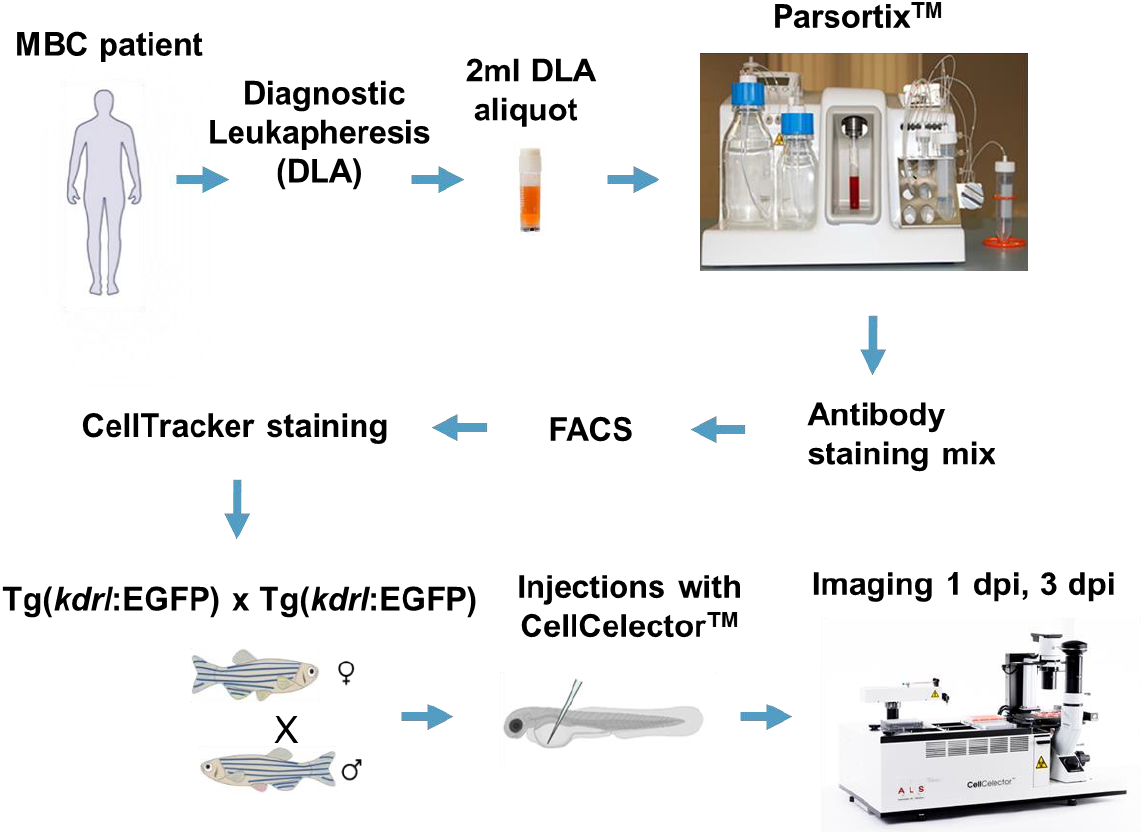
DanioCTC-Workflow. An MBC patient undergoes Diagnostic Leukapheresis (DLA). CTCs are enriched from a DLA aliquot with the Parsortix system and isolated by FACS after staining. Cells are further stained by CellTracker Red for tracking in zebrafish embryos. Single CTCs are isolated by using the CellCelector and injected into 2 dpf old Tg(*kdrl*:EGFP) zebrafish embryos. Adapted from (27, 28).

### Adaptations that were required to realize DanioCTC

#### Optimizing cell picking and injection

In order to determine the best suited capillary type for injecting the CTCs into the zebrafish larvae, the commercially available capillary of the CellCelector was compared with capillaries, which are prepared with a micropipette puller and regularly used for zebrafish injections (Figure 2). The self-prepared capillary is characterized by a long taper and a slightly bigger opening diameter (ca. 25 μm). The original CellCelector capillary, on the other hand, tapers quickly, is therefore rather short and its opening has a diameter of 20 μm. Since more cells stuck in the longer tapered tip of the self-prepared capillary, the use of this capillary type was discontinued and the CellCelector capillary was used. To improve the penetration of the CellCelector capillary into the zebrafish embryo, the tip was broken off with forceps increasing its diameter to approx. 25 μm. An angle of approximately 45° for the capillary opening proved to be the most effective for penetrating zebrafish tissue.

**Figure 2:**
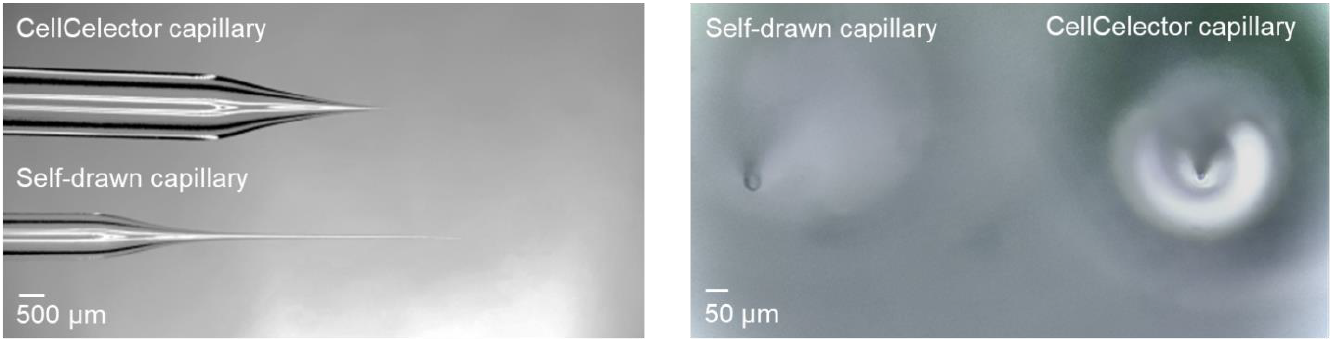
Capillaries. Self-prepared capillary for standard zebrafish embryo injections with a long taper narrowing and an opening diameter of approximately 25 μm. The CellCelector capillary tapers quickly, is rather short and has an opening diameter of 20 μm.

#### Adaptation of the CellCelector setup for cell injection

In order to enhance the functionality of the CellCelector, we collaborated with the manufacturers to modify the software. We programmed the joystick that allowed manual control of the robotic arm and injection process. Additionally, we created an automated injection process that allows for adaptation of the injection speed and volume. To simplify the workflow, we defined a fixed position for the zebrafish larvae and established a motion sequence for the robotic arm during injection.

#### Installation of a stereomicroscope at the CellCelector

One challenge in the injection procedure was to ensure that the entire process could be observed. As the CellCelector system uses an inverted microscope, it was not initially suitable for imaging from the top as required for injection into the DoC. To overcome this, we extended the system by installing a flexible stereomicroscope at the programmed deposition area of the CellCelector robotic arm. This allowed us to localize the DoC and position the capillary with precision. By aligning the CellCelector capillary with the DoC, we were able to achieve complete visualization of the injection process (as shown in Figure 3).

**Figure 3:**
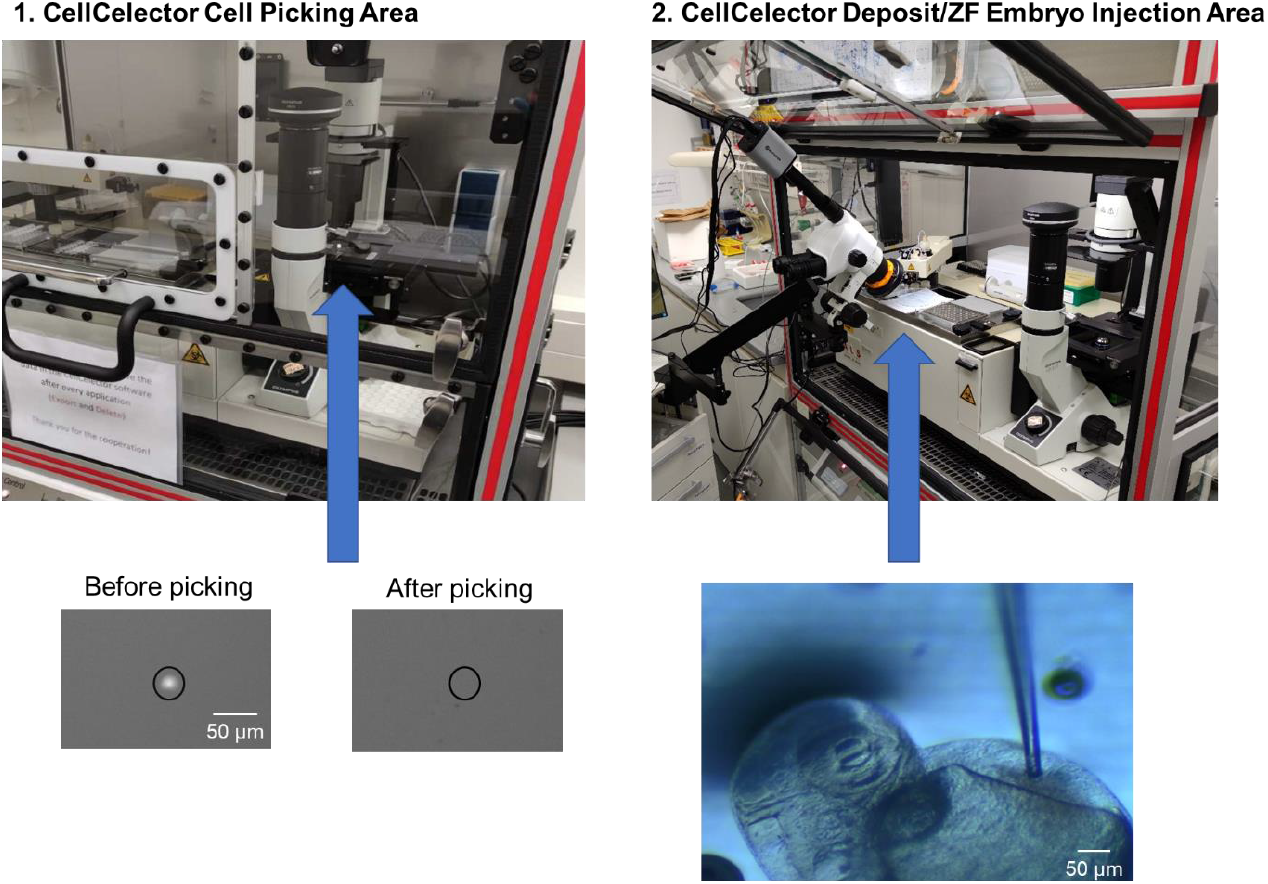
Adapted CellCelector setup: Representative images of the CellCelector picking area of picking a single CellTracker Red stained CTC and the attached stereomicroscope required for CTC injections in the deposit area of the CellCelector. 1. CellCelector cell picking area and image of region of interest before and after cell picking. 2. CellCelector deposit/zebrafish injection area and brightfield image of embryo being injected.

### Standard xenotransplantation workflow carried out with MDA-MB-231 cells

In our experiments, we used the standard workflow to inject GFP-labeled MDA-MB-231 cells into 2-day-old zebrafish embryos. A cell suspension of 1-2 μl was loaded into a glass capillary and 1-2 nl were injected into the DOC using the pneumatic PicoPump. The injected cells dispersed into various areas of the zebrafish embryos, including the head, trunk, and tail, which contains the caudal hematopoietic tissue, the site of blood formation at this larval stage (Figure 4). At 1 dpi, an average of 38.6 +/- 14.8 cells disseminated into the head region, 30 +/- 15.5 cells into the trunk region, and 123 +/- 31.4 cells into the tail region. However, by 3 dpi, only 30 +/- 20.5 cells were located in the head, 16.3 +/- 10.2 cells in the trunk, and 46.3 +/- 19.5 cells in the tail. This suggests that approximately 50% of the injected cells died and disappeared between 1 and 3 dpi, with more cells getting lost in the trunk and tail regions compared to the head region. Relative cell numbers can be found in Supplement Figure 2.

**Figure 4:**
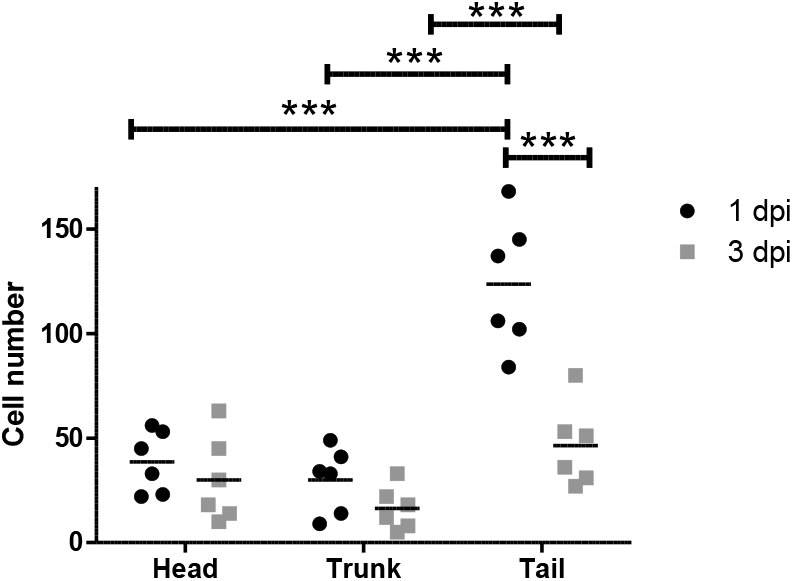
Dissemination of MDA-MB-231 cells after injection with the standard workflow into zebrafish larvae. Depicted are the absolute MDA-MB-231 cell dissemination into the head, trunk and tail at 1 and 3 dpi (n=6). dpi: days post injection. Repeated measured ANOVA test, ***0.0001 < p < 0.001

### Injection of MDA-MB-231 cells spiked into a DLA aliquot with the DanioCTC workflow

Since the CTCs will be isolated from DLA-products obtained from metastatic breast cancer patients, in the next step we mimicked such a sample by spiking about 5,000 MDA-MB-231 cells into a CTC-negative 2ml DLA aliquot (negative control) and processed it according to the workflow depicted in Figure 1 and Figure 5. Per embryo, (Tg(*kdrl*:EGFP)), 50 pre-labeled MDA-MB-231 cells were injected with the CellCelector, resulting in a total of 11 embryos. The zebrafish embryos were incubated and the injected cells were monitored at 1 and 3 dpi. The MDA-MB-231 cells disseminated to different areas in the head, trunk, and tail regions in varying numbers (Figure 5 A-D). On average, 3.5 +/- 3.2 cells disseminated into the head, 0.9 +/- 1.0 cells into the trunk, and 5.9 +/- 4.7 cells into the tail at 1 dpi (Figure 5 D). At 3 dpi, 3.8 +/- 2.0 cells were detected in the head, 0.8 +/- 1.1 in the trunk, and 3.9 +/- 4.4 in the tail, which is equivalent to approximately 83% of the cell numbers present at 1 dpi. The relative cell numbers are depicted in Supplement Figure 3. The MDA-MB-231 cells did not only accumulate but also partially extravasate from the blood circulation into surrounding tissue areas in the caudal hematopoietic tissue and other tail regions (Figure 5 C). This finding is consistent with the accepted metastatic nature of the MDA-MB-231 cell line (26, 29, 30).

**Figure 5:**
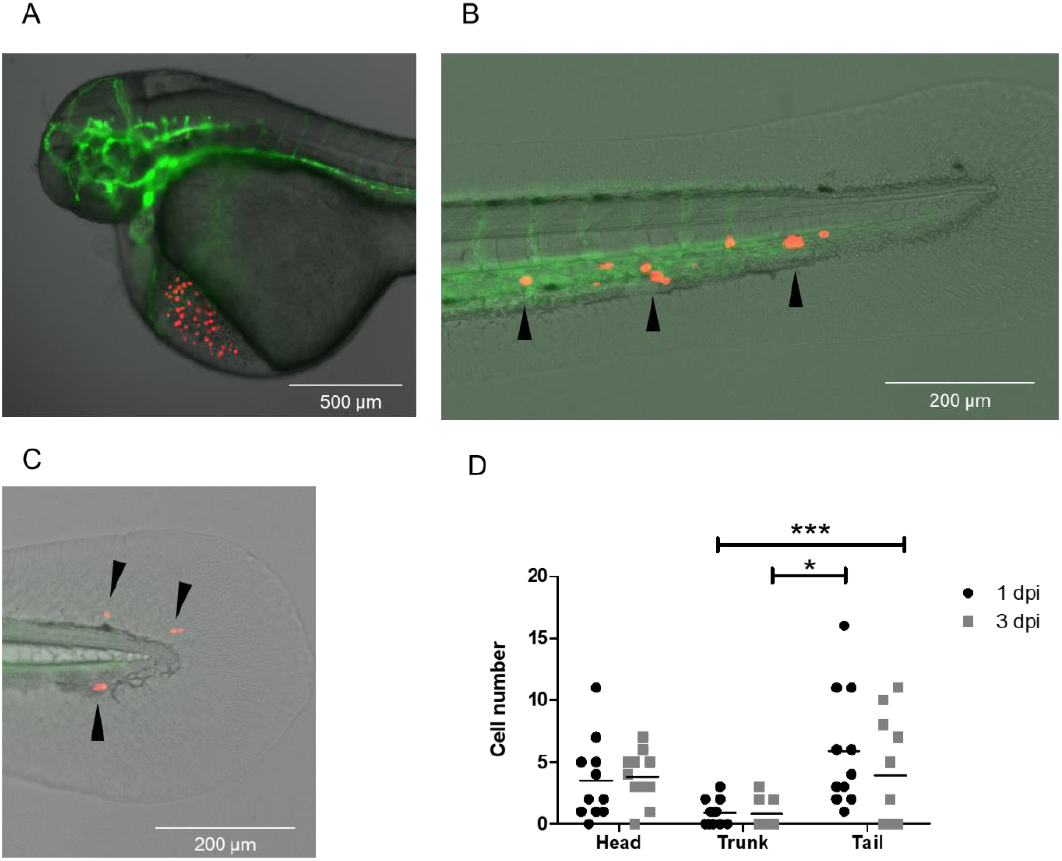
Dissemination of MDA-MB-231 cells spiked into a DLA sample after injection with the DanioCTC workflow. Cell localization was monitored at 1 and 3 dpi. A) Representative image of 50 MDA-MB-231 cells injected into the DoC of a zebrafish embryo (1 hpi). B) Representative image of the tail and CHT region at 3 dpi with localized MDA-MB-231 cells in the CHT (black arrowheads). C) Representative image of the tail region at 3 dpi showing extravasated MDA-MB-231 cells (black arrowheads). D) Absolute MDA-MB-231 cell numbers detected in the head, trunk and tail regions at 1 and 3 dpi (n=11 embryos). Green label: endothelium; red label: MDA-MB 231 cells. dpi: days post injection. Friedman test, *0.01 < p < 0.05; ***0.0001 < p < 0.001

### Injection of CTCs isolated from DLA aliquots with the DanioCTC workflow

A cryopreserved 2 ml DLA aliquot obtained from a patient with MBC was used for CTC enrichment using the Parsortix system, following our workflow (Figure 1). The resulting sample was labeled with an antibody cocktail for CTC isolation by flow cytometry. Isolated EpCAMpos/CD45neg cells were then stained with CellTracker Red to allow for tracking of CTCs in zebrafish embryos. Next, we transferred the sample onto a glass slide and used the CellCelector to pick 50 single CTCs semi-automatically, based on specific criteria: round, intact morphology with a cell diameter of 4–40 μm in brightfield (BF) and a positive CellTracker Red (TRITC) signal. We micromanipulated and injected each single CTC into the DOC of 2 dpf Tg(*kdrl*:EGFP) zebrafish embryos using the CellCelector. We repeated this process for a total of 50 CTCs per zebrafish embryo (n=9).

After injection, CTCs disseminated to different areas in the head, trunk, and tail (Figure 6), with an average of 0.4 +/- 0.7 CTCs in the head, 2.2 +/- 2.2 CTCs in the trunk, and 0.4 +/- 0.7 CTCs in the tail at 1 dpi. At 3 dpi, the mean number of CTCs detected in the head was 1.8 +/- 1.7, in the trunk was 0.6 +/- 1.0, and in the tail was 0.2 +/- 0.6, which is equivalent to approximately 87% of the cell numbers present at 1 dpi. We did not observe any extravasation events at 1 dpi or 3 dpi. See Supplement Figure 4 for relative cell numbers.

**Figure 6:**
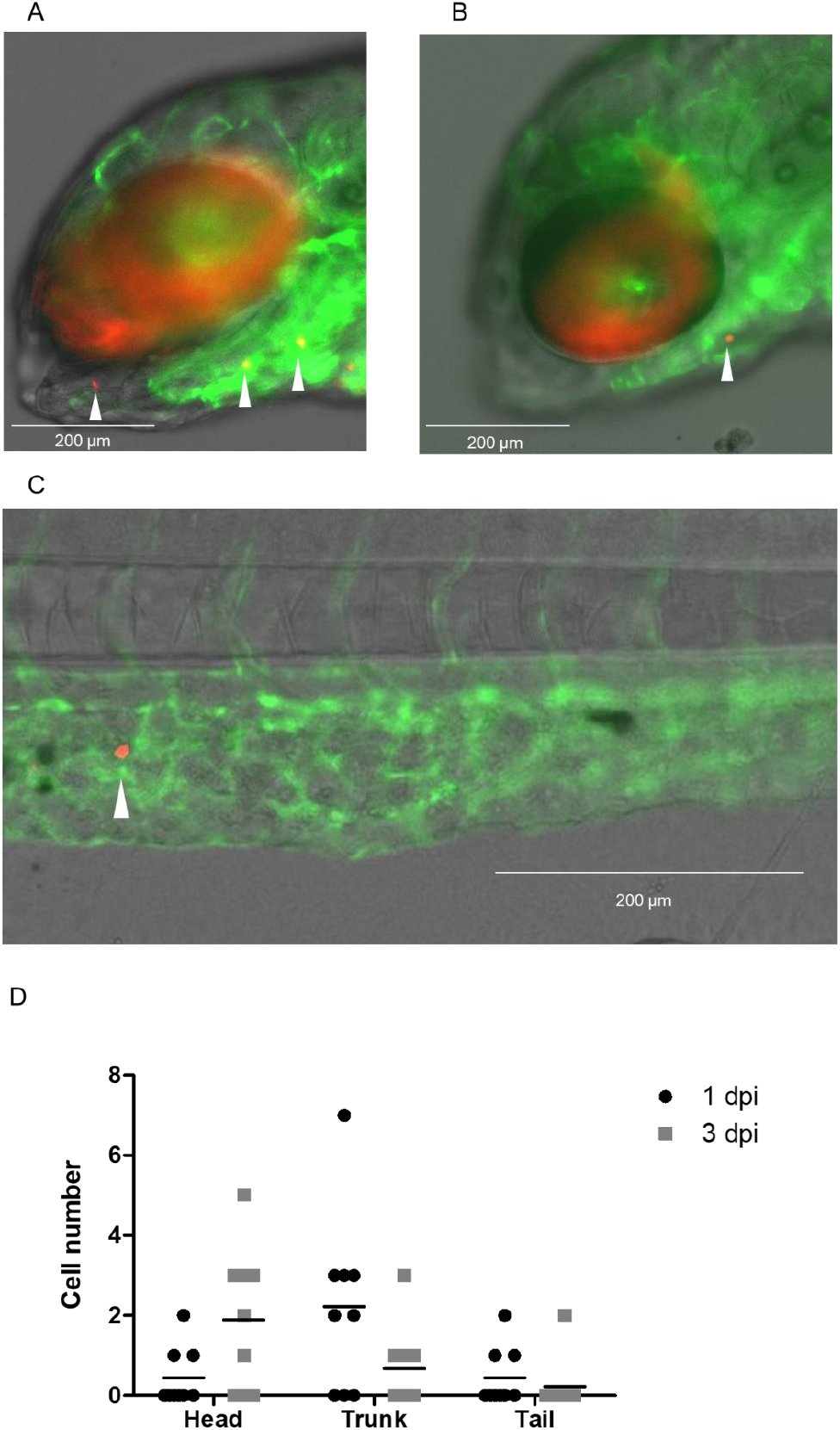
Dissemination of isolated CTCs of an MBC patient after injection with the DanioCTC workflow. Isolated CTCs, labeled in red, were monitored at 1 and 3 dpi and showed dissemination into the head, trunk and the tail. A) and B) Representative images of the head region of a Tg(*kdrl*:EGFP) positive embryo after injection of 50 CTCs cells (indicated by white arrowheads). C) Representative image at 3 dpi showing a disseminated CTC in the CHT of an embryo (indicated by white arrowhead). D) Absolute CTC numbers in the head, trunk and tail at 1 and 3 dpi (n=9). dpi: days post injection. Friedman test.

## DISCUSSION

Here we introduce DanioCTC, a seamless workflow that allows for the injection of low numbers of patient-derived CTCs into zebrafish embryos. This technique enables real-time studies of the differential metastatic traits of individual CTCs *in vivo*, despite their scarcity in cancer patients’ blood. Unlike previous methods which rely on high cell numbers and retrograde loading of cells into glass capillaries, our approach overcomes this technical limitation by integrating the injection process, viable CTC isolation, labelling, and detection into a reproducible workflow. With DanioCTC, we aim to preserve the competency of CTCs to metastasize by using directly isolated CTCs. Our innovative workflow combines the detection of CTCs from DLA products with *ex vivo* labelling techniques and direct injection using the automated micromanipulation system CellCelector, which we have modified and extended. DanioCTC facilitates the investigation of CTCs during the process of metastasis by enabling the injection of low numbers of patient-derived CTCs into zebrafish embryos. This approach provides a good basis for modelling the low frequency of CTCs typically found in cancer patients’ blood. The zebrafish embryo is an ideal *in vivo* model for studying CTC metastasis due to its transparency, fast development, high progeny number, and lack of adaptive immunity in initial life stages. In this transparent model, we can access the capacity of CTCs to arrest and adhere to the endothelium in a dynamic fashion, which may involve cell rolling and other processes that can be observed in tumor cells and immune cells found in the circulation (31). We can also study hemodynamic forces of blood flow that impact the arrest of CTCs. Compared to *in vitro* assays, which are somewhat static in nature and may not accurately reflect the low affinity and transient interactions of CTCs with endothelia, DanioCTC provides a more accurate way to monitor CTC transport and migration *in vivo*. In combination with other advantages (fast development, high progeny number, transparency, lack of adaptive immunity in initial life stages) zebrafish represents an ideal *in vivo* model to study CTC metastasis.

### MDA-MB-231 cell and CTC distribution in zebrafish larvae

We present the results of injecting MDA-MB-231 cells at both high and low cell numbers into zebrafish embryos, where they were frequently found in the head (including the CNS) and tail (CHT), which is consistent with the high incidence of brain and bone marrow metastases in triple negative breast cancer (32). The relative cell numbers in the head and tail changed over time, indicating a higher survival rate of MDA-MB-231 cells in the head. However, further investigations are needed to determine which individual tumor cells survive, as the redistribution of cells over time suggests variability in this aspect. MDA-MB-231 cells were observed to extravasate from blood vessels into the surrounding tissue, which is in agreement with previous findings (26, 29). In contrast, CTCs from a MBC patient injected into zebrafish larvae were primarily located in the head and trunk, unlike MDA-MB-231 cells that were present in the head and tail. The reasons for this are currently unclear. Flow conditions in the blood vessels of the respective domains, as well as the size of the cells, might influence the outcome. However, caution is warranted as this observation is based on a single experiment and DLA aliquot. Therefore, further investigation is required to determine if there is a correlation between brain metastases and localization to the head in zebrafish embryos. In the future, it will be important to isolate tumor cells from different regions of injected zebrafish embryos to compare their gene expression profiles with CTCs and DTCs from the patient to examine their expression profiles and clonal relationship.

### Advantages and limitations of DanioCTC

Our study, for the first time, employs a semi-automated single-cell micromanipulation system to demonstrate that CTCs from breast cancer patients can be directly injected into zebrafish embryos and monitored for several days through live imaging. While traditional mouse models have been utilized to mimic different stages of the metastatic cascade, zebrafish models offer a promising alternative. Previous research has showcased the potential of zebrafish models in investigating the biology of non-patient CTCs and CTC-clusters, indicating their utility in understanding the metastatic nature of cancer cells (20, 26, 29). For example, Berens et al. examined cancer cell extravasation in the zebrafish embryo tail 96 h after injection (29) with different cell lines. Zebrafish embryos injected with the MDA-MB-231 cell line exhibited the greatest number of extravasated cancer cells per embryo, which is in line with its increased metastatic nature. In agreement with this, Asokan et al. showed that MDA-MB-231 cells were capable of invading avascular tail fin tissue while normal breast epithelial cells were not (26). At 4 dpi, 31.8 % of injected MDA-MB-231 cells had extravasated into the caudal tail. Martinez-Pena and colleagues revealed that CTC-clusters from MDA-MB-231 cells disseminated at a lower frequency than single MDA-MB-231 CTCs in the zebrafish, showed a reduced capacity to invade but had a higher survival and proliferation capacity in the temporal follow-up than single CTCs (20).

Despite those advancements, cell lines as ‘surrogate CTCs’ have been used in all previous studies due to limitations in the injection process to investigate the differential behaviour of single CTCs and CTC-clusters in zebrafish embryos. It is important to note that subtle differences have been observed between cell line-derived surrogate CTCs and patient-derived CTCs (33-35) and caution should be exercised when interpreting the results.

We provide a proof-of-concept for the DanioCTC workflow that allows us to inject isolated CTCs from a single MBC patient into zebrafish embryos, to the best of our knowledge, for the first time. This case study, along with the established setup, will allow for further experiments on isolated CTCs, accompanied by ongoing improvements in the equipment and software. Future enhancements in the procedure and setup will reduce the processing time and increase the applicability of the method in personalized medicine approaches.

In summary, our study establishes an innovative workflow for the injection of MBC-derived isolated CTCs into zebrafish embryos, enabling live imaging of patient-derived CTC dissemination in xenografts over time. This approach may become a valuable tool for personalized medicine and in-depth exploration of CTC metastatic biology, given its capability to work with only a few dozen cells per embryo.

## ACKNOWLEDGEMENTS

We thank Katrin Lambert and Anastasia Kurzyukova for help in establishing the microinjection procedure, Manja Wobus and Martin Bornhäuser for providing cells, the Light Microscopy core facility at the CMCB for their support, and Marika Fischer, Silvio Kunadt and Daniela Moegel for excellent fish care. We thank Benjamin Odermatt and the Zebrafish Core-Facility at the Medical Faculty Bonn.

## FUNDING AND CONFLICT OF INTEREST

This work was supported by the German Research Foundation (DFG) Transregio 67 (project 387653785), the DFG SPP 2084 μBone (project KN 1102/2-1 to FK and project HO 1875/26-1 employing FR to FK). The work at the TU Dresden was co-financed with tax revenues based on the budget agreed by the Saxonian Parliament (‘Landtag’). The work in Duesseldorf was partly funded by the German Cancer Foundation (DKH) Priority Program ‘Translational Oncology’ (Grant 70112504).

## Notes

**Conflict of interest:** The authors declare no potential conflicts of interest.

### Competing Interest Statement

The authors have declared no competing interest.

