## Supplement for "DanioCTC: Injection of circulating tumor cells from metastatic breast cancer patients in zebrafish xenografts for analysis of metastasis"

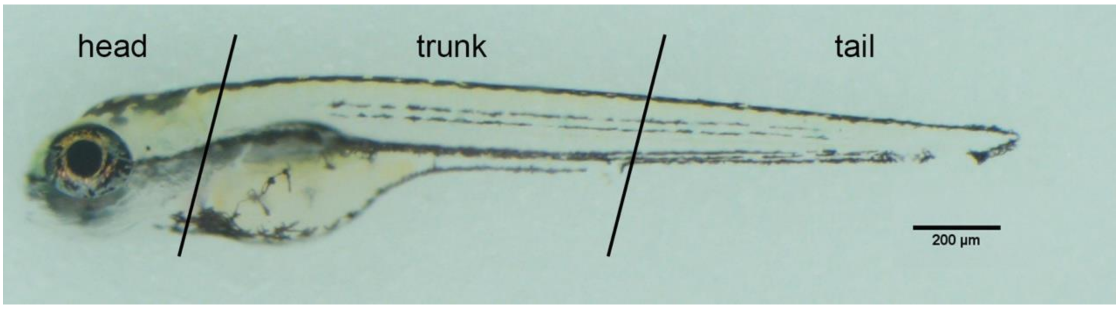


**Supplement Figure 1: Regions of the zebrafish larva:** The larval body was divided into three regions head, trunk, and tail for cell count analysis.

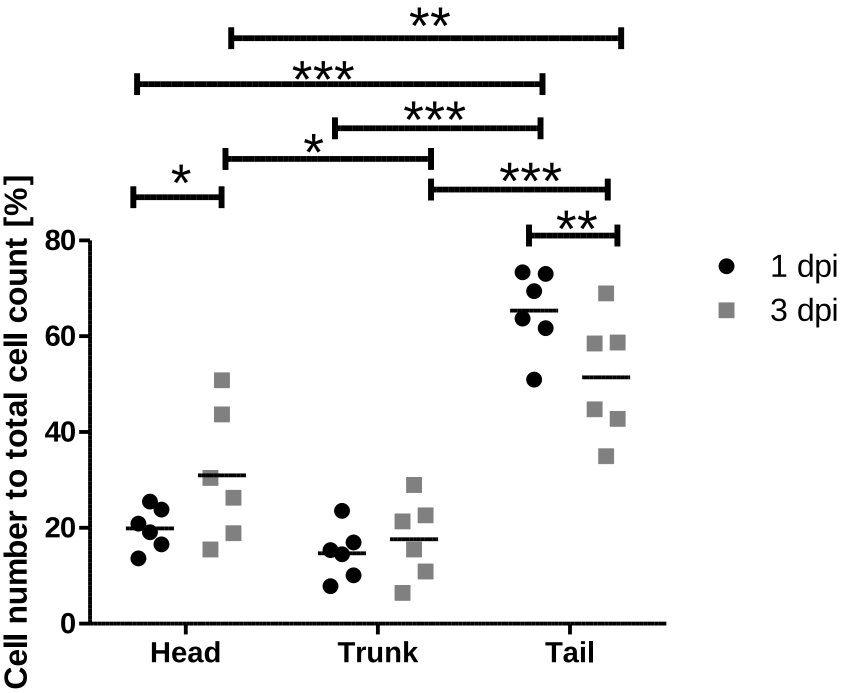


**Supplement Figure 2: Dissemination of MDA-MB-231 cells after injection with the standard workflow into 2dpf zebrafish larvae**. Depicted are the relative numbers of

MDA-MB-231 cells localized in the head, trunk and tail regions at 1 and 3 dpi (n=6). 19.9 % of the cells were located in the head, 14.7 % of cells in the trunk and 65.3 % in the tail at 1 dpi. At 3 dpi, 30.9 % of the cells were located in the head region, 17.6 % in the trunk and 51.4 % in the tail. dpi: days post injection. dpi: days post injection, Repeated measured ANOVA test, *0.01 < p < 0.05; **0.001 < p < 0.01; ***0.0001 < p < 0.001

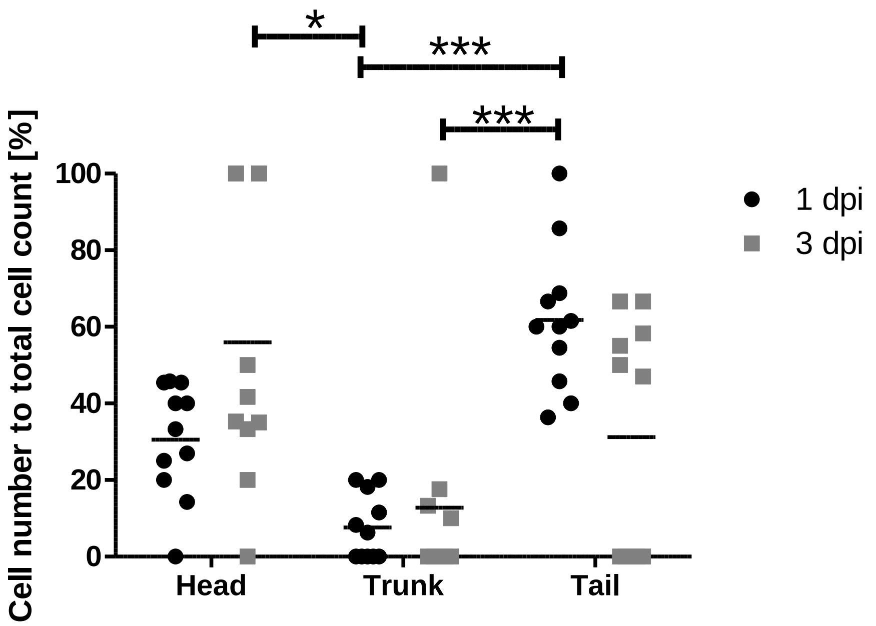


**Supplement Figure 3: Dissemination of MDA-MB-231 cells spiked into DLA samples after injection with DanioCTC workflow into zebrafish larvae.** Depicted are the relative numbers of MDA-MB-231 cells localized in the head, trunk and tail regions at 1 and 3 dpi (n=11). 30.5 % of the cells were located in the head, 7.6 % of cells in the trunk and 61.7 % in the tail at 1 dpi. At 3 dpi, 55.9 % of injected cells were detected in the head, 12.8 % in the trunk and 31.2 % in the tail. dpi: days post injection, Friedman test, *0.01 < p < 0.05; ***0.0001 < p < 0.001

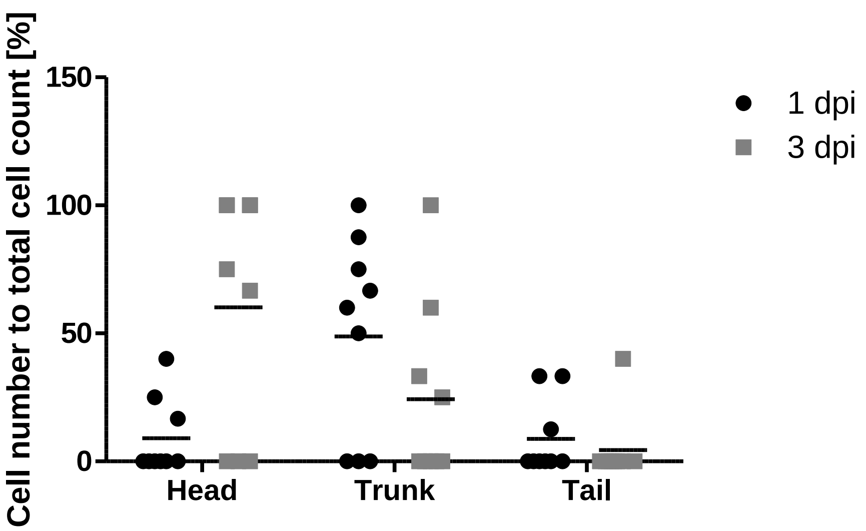


**Supplement Figure 4. Dissemination of isolated CTCs of an MBC patient after injection with DanioCTC workflow into zebrafish larvae.** Depicted is the relative CTC dissemination into the head, trunk and tail at 1 and 3 dpi (n=9). 9% of the CTCs were present in the head, 48,7 % in the trunk and 8,7% in the tail at 1 dpi. At 3dpi, their numbers corresponded to 60.1%, 24.2%, 4.4% in the head, trunk and tail, respectively. dpi: days post injection. Friedman test.
